# A new oculomotor model demystifies “Remarkable Saccades”

**DOI:** 10.1101/2024.06.14.599100

**Authors:** Heinen, Chandna, Singh, Watamaniuk

## Abstract

Hering’s Law of binocular eye movement control guides most oculomotor research and supports diagnosis and treatment of clinical eye misalignment (strabismus). The law states that all eye movements are controlled by a unitary conjugate signal and a unitary vergence signal that sum. Recent evidence of temporally asynchronous inter-eye rotations during vergence (Chandna et al., 2021) challenges the viability of a unitary vergence signal. An alternative theory that might explain these anomalous results posits that the eyes are controlled independently. Yet independent control fails to explain a phenomenon known as “Remarkable Saccades” where an inappropriate saccade occurs from an eye aligned on a target during asymmetric vergence (Enright, 1992). We introduce a new model formulated to describe the Chandna et al. (2021) midline vergence result that generates remarkable saccades as an emergent property. The Hybrid Binocular Control model incorporates independent controllers for each eye with a cortical origin that interact with a unitary conjugate controller residing in brainstem. The model also accounts for behavioral variations in remarkable saccades when observers attend to an eye. Furthermore, it suggests more generally how the eyes are controlled during vergence and other voluntary eye movements, thus challenging documented oculomotor neural circuitry and suggesting that refinements are needed for clinical oculomotor interventions.

## Introduction

Hering’s Law of binocular eye movement control guides virtually all oculomotor research and is the basis for diagnosis and intervention of clinical eye misalignment known as strabismus. The primary law states that the eyes are yoked and move identically together as a pair (Hering, 1868). A complementary law is posited for viewing targets in depth, which states that a unitary command rotates the eyes oppositely and symmetrically to either converge or diverge (Hering, 1868). Because eye movements are thought to be guided by these unitary commands, eye movement scientists who display stimuli on a tangent screen typically measure the rotation of one eye, assuming that the second eye behaves identically. When a target is displaced in depth, the assumption is one eye’s motion is a mirror image of the other’s, and clinical practitioners generally observe one eye and assume a covered eye behaves according to Hering’s Law.

Oculomotor science has enjoyed an incredibly prolific tenure. This success is at least partially attributable to the fact that oculomotor systems are described by sophisticated models. These are generally dynamic systems models that generate continuous output from continuous input. The models also incorporate feedback, which is a common feature of biological systems. The saccadic eye movement system, which rapidly moves the eyes from one visual target to another, has been modeled more extensively than any other oculomotor system. It is characterized as receiving an error signal from the retina about the location of a visual target. This error is passed to saccadic circuitry that corrects the error via a feedback loop that is closed on the retina (Robinson, 1973). The smooth pursuit system, which follows moving objects, is also modeled as a feedback system. This system minimizes retinal velocity error between the eyes and a moving target (Robinson et al., 1986; Krauzlis & Lisberger, 1989). The vergence system moves the eyes between objects that are located at different depths. Although much less studied than pursuit or saccades, vergence models similarly use feedback to minimize retinal disparity error that arises when an object is displaced in depth (Hung et al., 1986).

Because all these models assume Hering’s Law, they accept a single “retinal error” input, and generate a single eye rotation command resulting from a unitary signal that controls both eyes. However, many eye movements made in natural scenes shift gaze between objects that have a depth component (i.e., they do not lie on the horopter), and hence purely conjugate eye movements are inappropriate to divert gaze to them. Nor are these eye movements strictly on the midline, where a unitary vergence command theoretically drives the eyes with exactly opposite rotations. To handle asymmetric eye rotations, Hering postulated that conjugate and vergence eye movement signals simply sum, and as a result, a unitary command still is sufficient to drive the eyes to any location in visible space (**Fig 1A**).

**Fig 1.**
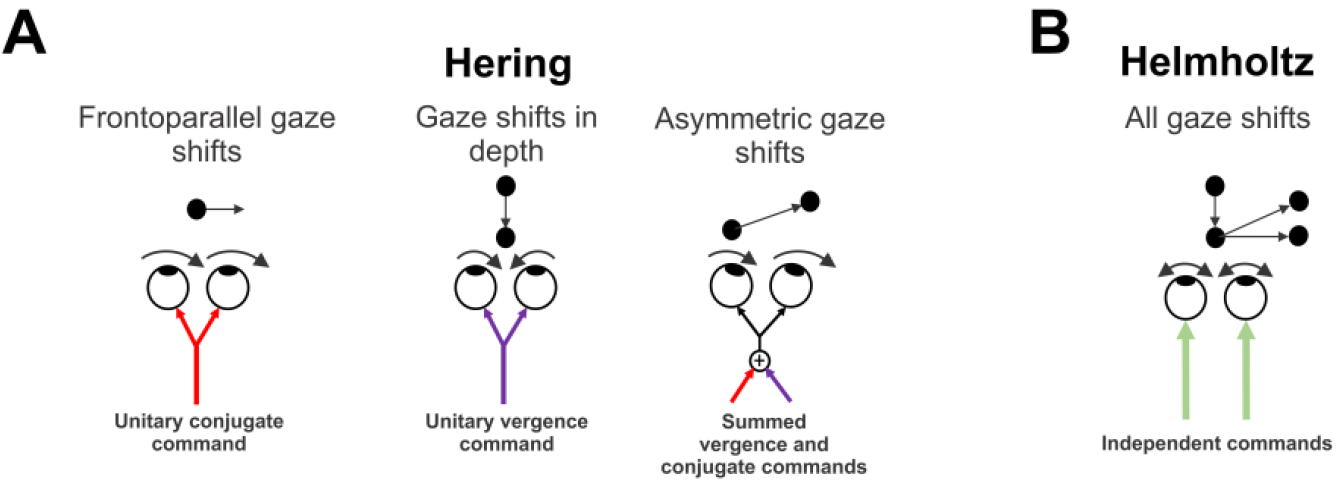
Classic models of oculomotor control. A) Hering’s law of yoked eye movement control. B) Helmholtz’s law of independent control.

Recent work casts doubt on the existence of a unitary vergence command. Chandna et al. (2021) occluded an eye in normal observers as they used smooth pursuit to make vergence movements to targets moving on the midline. Interestingly, the rotation of the covered eye was completely unpredictable from the viewing eye’s rotation, or from the motion of the target. While the viewing eye rotated appropriately to follow the target’s motion in depth consistent with vergence, the covered eye did not. It sometimes rotated with a weak vergence component, but more often it had either negligible movement or rotated conjugately with respect to the viewing eye. Furthermore, the covered eye’s rotation was delayed relative to the viewing eye’s rotation, sometimes on the order of seconds. The delay of the covered eye’s rotation is incompatible with a unitary vergence command driving the eyes (for discussion, see Chandna et al., 2021).

However, an alternative theory proposed by Helmholtz (1867) could account for the Chandna et al. (2021) results. Helmholtz theorized that the eyes are controlled independently, and that we learn to move them together (**Fig 1B**). The independent control theory was recently revived given evidence of single-eye specific signals being present in brainstem structures previously thought only to generate conjugate eye movements (Zhou & King, 1998). To capture these results, these researchers implemented independent control in a computational model (King & Zhou, 2002). However, while theoretical independent control of the eyes ostensibly accounts for any eye movements including the Chandna et al. (2021) results, it too has problems. The most basic problem is that an occluded eye that lacks visual input should not move regardless of the viewing eye’s rotation. Yet, the occluded eye in Chandna et al. (2021) usually continued to rotate and did so in an unpredictable fashion.

Another perhaps more daunting challenge for independent control is a curious oculomotor phenomenon known as “The Remarkable Saccades of Asymmetrical Vergence” (Enright, 1992). Remarkable saccades are such named because they are inappropriate saccades generated when stimuli are arranged using the Müller paradigm, in which two targets are aligned on the optical axis of one eye at different distances. Remarkable saccades occur when observers generate saccades between the two targets. For example, during convergence, when a saccade is made from the far to the near target, the unaligned eye executes the saccade as it should. However, the aligned eye, which should remain stationary since the targets are aligned to it, generates a smaller inappropriate saccade in the same direction as that executed by the unaligned eye (away from both targets). Since the visual input specifies that the aligned eye should not move, an independent control architecture should hold the aligned eye stationary.

Here we present a model that was constructed to account for the Chandna et al. (2021) data but that generates remarkable saccades as an emergent property. A key aspect of the model is that it accepts separate left and right eye inputs and issues separate left and right eye commands. This is different from virtually all other oculomotor models, except those explicitly designed for independent control (King & Zhou, 2002). Most models of conjugate eye movements or vergence are cyclopean in that they accept a single retinal input and issue a single command. At its core, our model has a rapid conjugate subsystem that generates saccades in the two eyes with the same vector, as well as two independent controllers, one for each eye, that generate slower eye movements. These independent controllers interact with the output of the conjugate system to generate asymmetric eye movements. We call our model Hybrid Binocular Control (HBC) model. Here we test it using data from the Enright (1992) paper that investigated remarkable saccades.

### Model Structure

The model was constructed using Simulink, a component of the Matlab software suite (The MathWorks Inc., Natick MA). Simulink enables construction of dynamic systems models that emulate electronic circuits, and its elements are largely based upon analog electronic components.

**Fig 2** shows a schematic circuit of the HBC model. The left and right eyes send input to their respective perceptions. Here perception is implemented as a simple summation between individual eye inputs and their respective feedback loop. Note that the details of the perceptions will need to be determined by future experiments. However, for the purpose of the present simulations, we simplify and equate perceptions to the retinal input. The virtual target is the interaction of left and right eye percepts which yields a unified percept and was proposed to account for the results obtained under monocular viewing reported in Chandna et al. (2021). In the center is a rapid conjugate subsystem. The subsystem has a right-brain conjugate component that directs both eyes leftward and a left-brain component that directs both eyes rightward. The controllers have identical transfer functions, with short time constants (*τ*=0.5 sec) that drive the eyes rapidly, consistent with rapid saccade generation. While each controller receives input from its respective eye, the output passes through a single node which in isolation would drive the eyes identically, but within our model the eyes are also influenced by the independent controllers. Mutual inhibition is implemented between the conjugate controllers so that simultaneously triggered but oppositely-directed saccades (e.g., vergence saccades) cancel in which case the independent controllers drive the eye movements. Gain values are shown in **Table 1**.

**Table 1.** Model parameters for typical remarkable saccades. Parameters for subject AB from Enright (1992) (shown in Fig 5). Left and right eye gain values (G_l_ and G_R_) assume the right eye is aligned with both targets.

|  |  |
| --- | --- |
| left eye gain ($G_l$) | 2 |
| right eye gain ($G_r$) | 1 |
| left eye independent gain ($G_{lR}$) | 110 |
| right eye independent gain ($G_{lL}$) | 110 |
| conjugate gain ($G_c$) | 40 |
| attention multiplier ($G_a$) | 1 |

**Fig 2.**
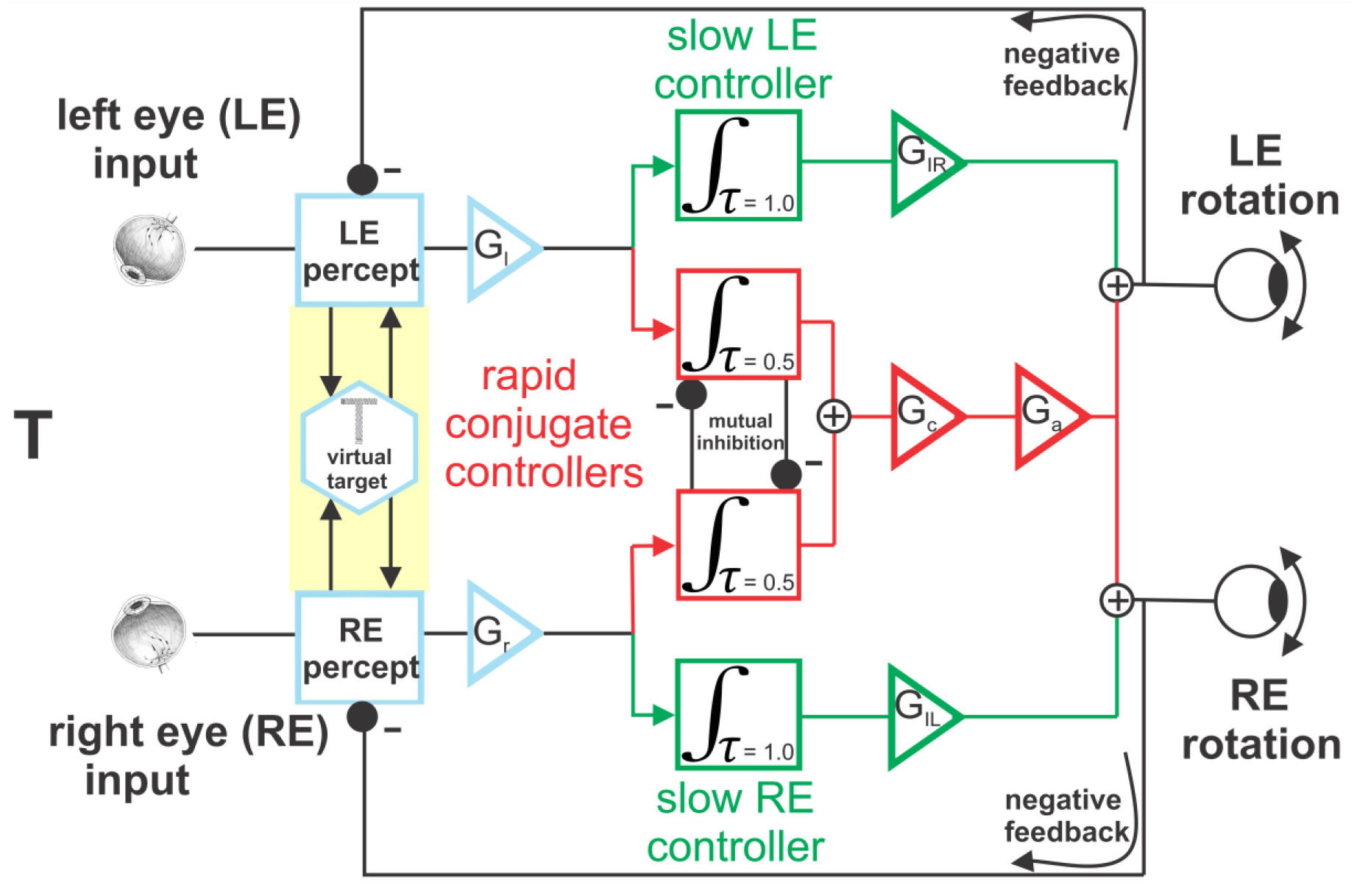
The Hybrid Binocular Control model. Separate eye inputs feed perception (blue) which generates the binocular virtual target (suggested by monocular occlusion). Independent controllers (green) slowly modulate each eye’s rotation, assist in generating asymmetric eye movements and may reside in cortex. Conjugate controllers (red) rapidly rotate the eyes as a pair, confer saccades their high speed and may reside in the brainstem. The conjugate controllers mutually inhibit each other to cancel simultaneous oppositely-directed saccades (on midline). Note that percept and virtual target components are conceptual, and not implemented computationally. Note also that negative feedback minimizes error of the perceived target’s position, unlike prevailing models that minimize retinal position error.

The model also implements independent control of each eye. The independent controllers, which are identical here, have their own transfer functions with longer time constants (*τ* =1.0 sec) (see **Table 1**). These controllers generate a more sluggish eye movement than does the conjugate subsystem, consistent with smooth, slow oculomotor behaviors like pursuit and vergence. Each independent signal has a separate gain (GIL & GIR), which here are set to the same value, and the conjugate signal has its own, singular gain (Gc) (see **Table 1**). The independent controllers each interact with the conjugate controller via an additive connection before the individual eye rotation commands are issued. Thus in our model, Hering’s unitary vergence command is replaced by commands issued by an independent controller for each eye.

A common feature of oculomotor models is dynamic feedback that minimizes either error between the location of an image on the retina and the fovea, or the error between the velocity of a retinal image and the eyes’ rotational velocity. However, our model does not simply minimize retinal error, rather it minimizes the error between the eye and the *perceived* target. Therefore, the feedback loop closes on perception, as suggested by other results where eye movements are directed to perceived attributes of the retinal stimulus rather than to a retinal error signal (e.g., Beutter & Stone, 1998; Heinen & Watamaniuk, 1998; Watamaniuk & Heinen, 1999; Madelain & Krauzlis, 2003). For simplicity, in tests of the model here we assume that perception is veridical with the retinal signal.

### Tests of the Model

#### Vergence pursuit

First, we present data from midline smooth-pursuit vergence (Chandna et al, 2021) that was fit by a computational instantiation of our conceptual model proposed in that same paper. **Fig 3A** shows vergence movements from Chandna et al. (2021) for one observer during binocular viewing, with fits from our model superimposed (green traces). By adding only a delay to the perceptual component of the model driving the covered eye, we simulated the asynchronous and reduced amplitude movements of the covered eye (**Fig 3B**). The delay could arise because the covered eye controller must readout a weak signal from the virtual target that is normally strengthened by direct visual input. For the covered eye controller to activate, more time may be necessary to accumulate sufficient neural activity at synaptic junctions to reach threshold for initiating an action potential (Kandel et al., 2000).

**Fig 3.**
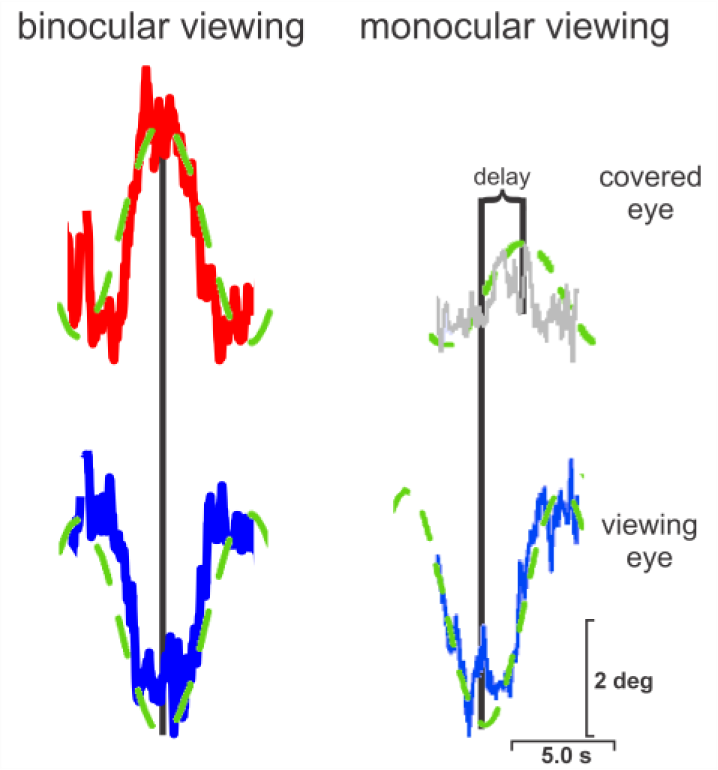
Vergence pursuit and model fits from Chandna et al. (2021). Shown are single cycles of sinusoidal smooth pursuit on the midline during binocular or monocular viewing. Behavioral data shown as red (right eye), blue (left eye) or grey (covered right eye) curves. Model fits (green dashed curves) capture binocular viewing, and delay in covered eye.

#### Vergence saccades

In our model, conjugate controllers were incorporated to explain conjugate intrusions in the monocular vergence data from Chandna et al. (2021). These controllers generate either rightward or leftward conjugate eye movements. Rotation of the eyes in opposite directions during vergence would cause both conjugate controllers to be activated simultaneously. To prevent instability in the model, we instantiated mutual inhibition between them, an arrangement supported by mutual inhibition between inhibitory burst neurons (IBNs) in the brainstem saccade circuit (Shinoda et al., 2008). Interestingly, this arrangement allows our model to generate vergence saccades, during which eye velocity is much lower than during conjugate saccades (**Fig 4**), a phenomenon observed in primates (Quinet et al., 2020).

**Fig 4.**
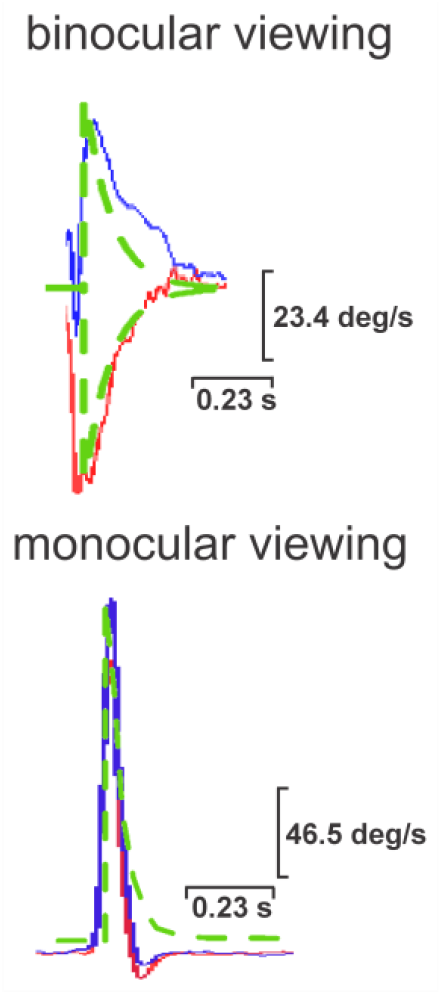
Vergence saccades and model fits. Model fits of eye velocity during binocular vergence saccade, and monocular, conjugate vergence saccade. Data collected from one observer in our laboratory. Details as in Fig 3. Note that in the bottom plot the blue and red curves overlap.

#### Remarkable saccades – an emergent feature

While the model was developed to explain smooth pursuit vergence on the midline during binocular and monocular viewing, as an emergent property it also generates the remarkable saccades of asymmetric vergence (Enright, 1992). Enright discovered the remarkable saccades using the “Müller paradigm” in which two targets are aligned at different distances along the optical axis of one eye. In that study, he instructed observers to make verbally-cued saccades between the targets to produce asymmetric vergence. During asymmetric vergence in the Müller paradigm, no movement should occur in the aligned eye as one target is directly behind the other. Yet the aligned eye is curiously deflected away from the target in the same direction as the unaligned eye.

Figure 5. shows raw position traces from the left and right eyes from a representative observer generating a remarkable saccade (from Enright 1992, Figure 3), with fits of our model superimposed. The left eye was aligned on two targets that were positioned 18 and 70 cm from the observer. The observer shifted gaze from the far to the near target to enact an asymmetric vergence saccade. Here it can be seen that the unaligned right eye (bottom panel) made a reasonable saccade from the far to near target, with an amplitude of approximately 9 deg. While the left, aligned eye, inappropriately made a saccade in the same direction as the right eye with a magnitude of ∼3 deg (top panel). The structure of our model allows it to capture the characteristics of both the left and right eyes during remarkable saccades (blue and red lines). Appropriate remarkable saccade magnitude was accomplished merely by adjusting model gain parameters. It is important to note that if the eyes were independently controlled by retinal signals (e.g., Zhou & King,1998), the aligned eye should remain stationary.

All observers in Enright’s (1992) experiment generated the remarkable saccades, i.e., saccades in the aligned eye that were in the same direction as those in the unaligned eye.

**Fig 5.**
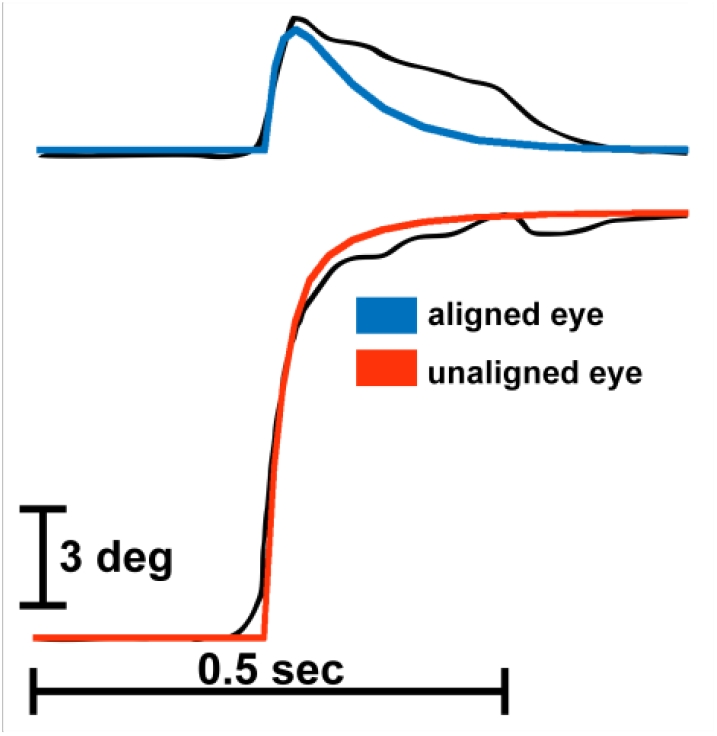
Model fit of remarkable saccade traces. Shown are raw eye position traces from a single trial (black traces) with model fits superimposed (blue, aligned eye and red, unaligned eye traces). Eye traces adapted from Enright (1992).

However, the size of the unaligned eye saccade, and even the size of the aligned eye saccade was highly variable between subjects, as well as within subjects from trial-to-trial. While our model does not have noise that would introduce trial-to-trial variability, it does generate remarkable saccades with a size that is a function of the magnitude of the saccade made in the aligned eye (**Fig 6**) as reported by Enright (1992; his Fig 6).

**Fig 6.**
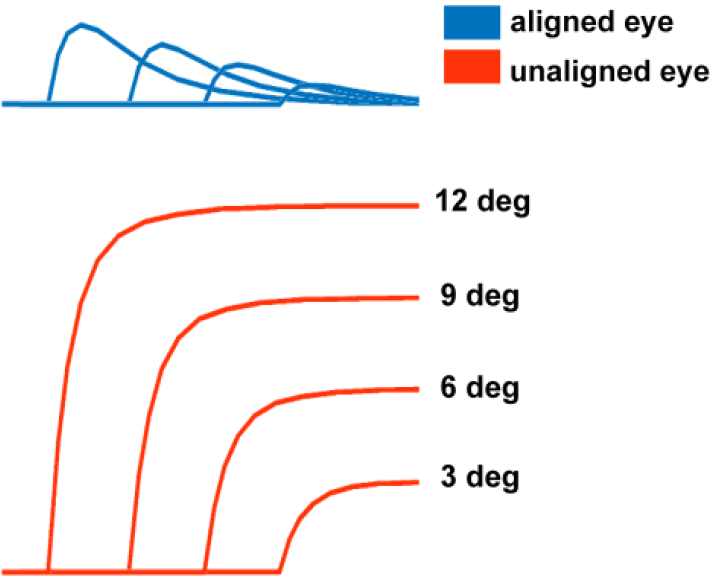
Remarkable saccades are a function of aligned eye saccades. At top are a family of eye position traces generated by the model for the aligned eye for saccade target separations ranging from 3 to 12 deg. At bottom are accompanying unaligned eye traces.

Enright (1992) quantified the remarkable saccades by computing the ratio of the magnitude of the saccade in the unaligned eye to that of the saccade in the aligned eye (UE /AE). While there was substantial variability in the ratios between subjects (and within each subject) with the target displacement used in the experiments (18 and 70 cm), our model replicated these ratios for the different subjects when we adjusted the value of just a single parameter, the output gain on the conjugate controller **(Gc)** (see **Fig 2**). **Fig 7A** shows the UE/AE ratios for subject AB from Enright (1992), along with the model’s performance. The conjugate controller gains for model fits to individual subject data are shown in **Table 2**. Note that even within a subject, the UE/AE ratios were highly variable. We determined variability for our model fits in Fig. 7 by generating unaligned eye saccades to 30 randomly selected target offsets between 12-16 degrees, corresponding to the expected movement of the unaligned eye based on Enright’s (1992) observations. Divergence ratios were similar to convergence ratios, and **Fig 7C** shows both convergence and divergence ratios from subject HP as well as model performance. Besides the UE/AE ratio, the difference in saccade magnitudes was also quantified in Enright, and **Fig 7B** shows that saccade amplitude differences (unaligned – aligned eye saccade magnitudes) overlapped for both subject AB and the model.

**Table 2.**
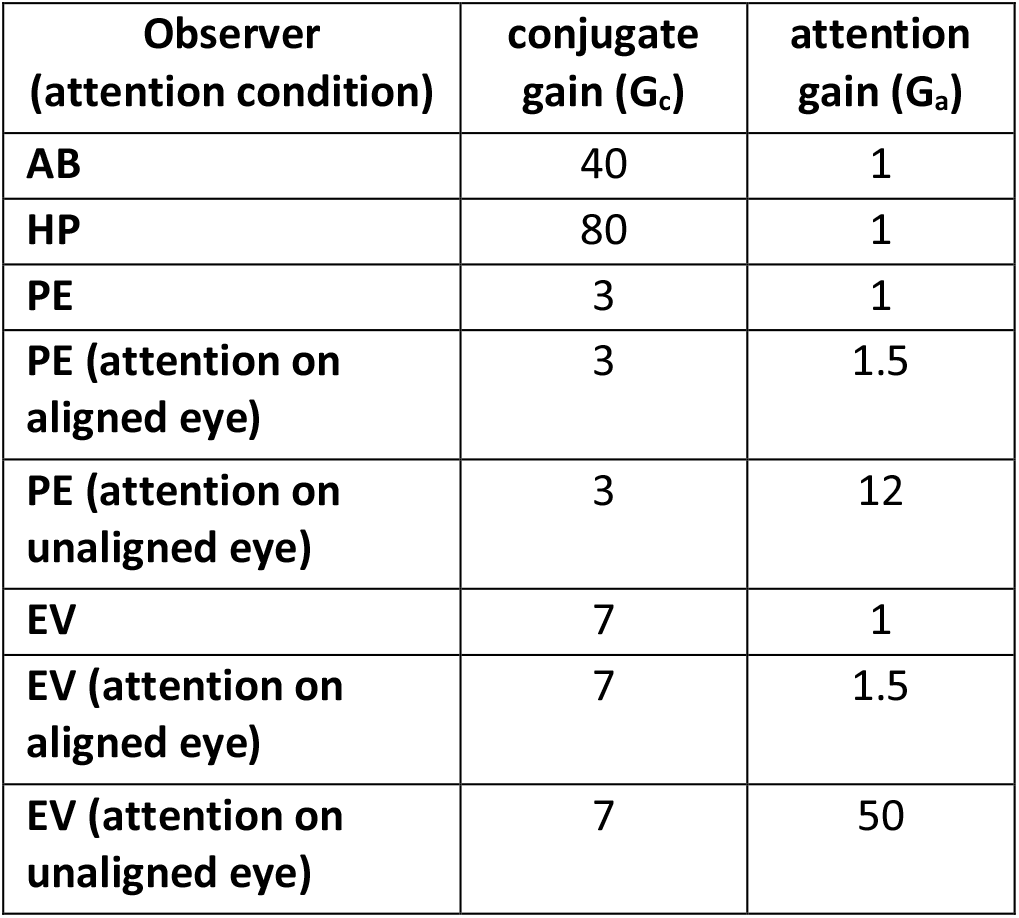
Gain values for different subjects and conditions.

| Observer<br>(attention condition) | conjugate<br>gain ($G_c$) | attention<br>gain ($G_a$) |
| --- | --- | --- |
| AB | 40 | 1 |
| HP | 80 | 1 |
| PE | 3 | 1 |
| PE (attention on<br>aligned eye) | 3 | 1.5 |
| PE (attention on<br>unaligned eye) | 3 | 12 |
| EV | 7 | 1 |
| EV (attention on<br>aligned eye) | 7 | 1.5 |
| EV (attention on<br>unaligned eye) | 7 | 50 |

**Figure 7.**
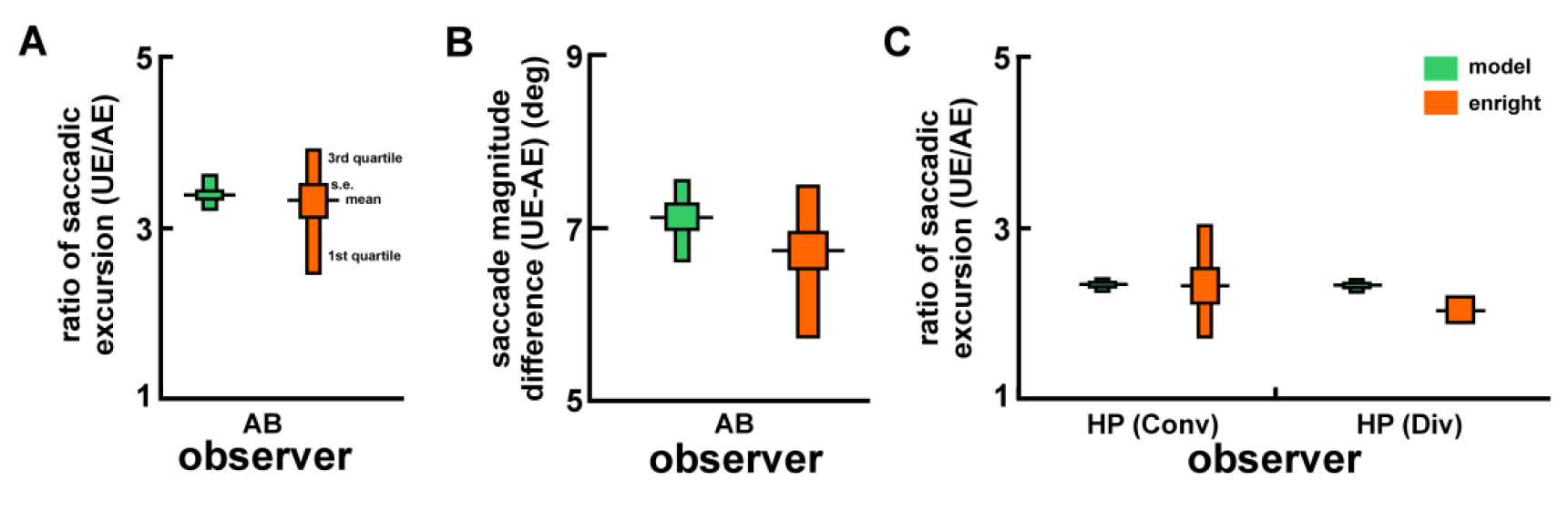
The model replicates the remarkable saccade for individual subjects. A) Ratios of unaligned saccade magnitude to aligned saccade magnitude (UE/AE) for subject AB. B) The difference in absolute saccade magnitude between the unaligned and aligned eye for subject AB. C) Saccade ratios for convergence (Conv) and divergence saccades (Div) for subject HP. On each datum, black horizontal lines indicate the mean, wide box indicates ±1 standard error, and the narrow box extends from the 1^st^ to 3^rd^ quartile, similar to Enright (1992).

Cognitive factors might have contributed to the different ratios of saccade amplitudes seen both within and between observers in Enright’s (1992) remarkable saccade experiment. Attention is one such factor, a well-known modulator of behavior, and Enright assessed its contribution to remarkable saccades. Enright had subjects make saccades to targets on the Müller axis while attending either to the eye that was aligned to the targets, or to the unaligned eye. While attending to the aligned eye did not affect performance, attending to the unaligned eye increased the UE/AE ratio.

Interestingly our model matched the human attention modulation (**Fig 8**) merely by changing the attentional gain on the conjugate controller (GA) (see **Fig 2**). **Table 2** shows the values of GA used to obtain the attention results.

**Figure 8.**
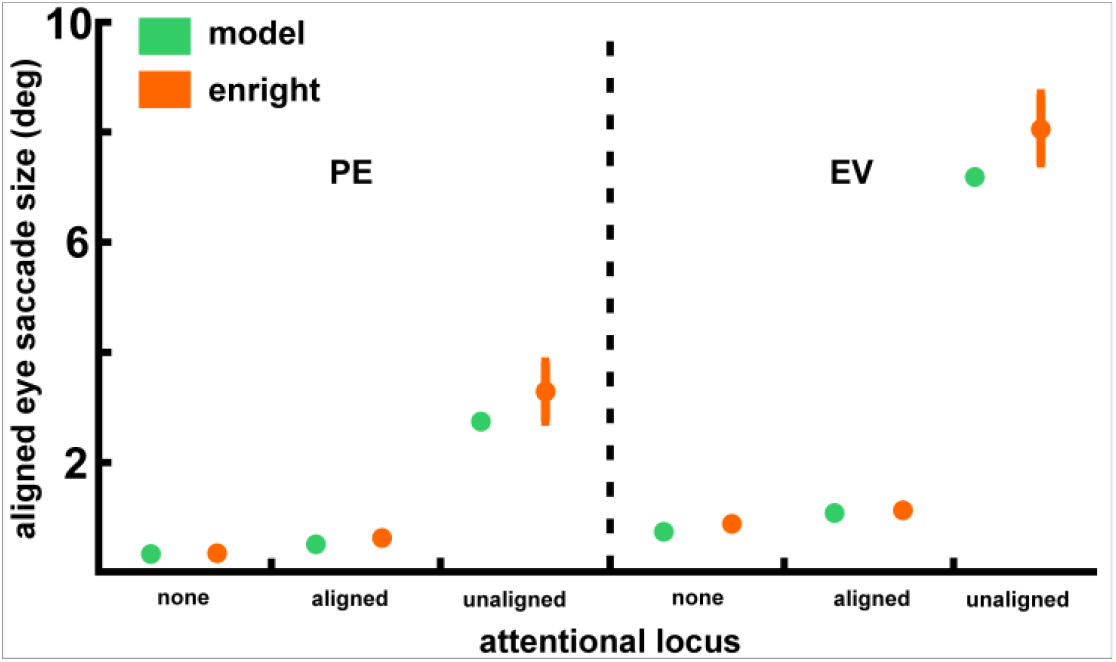
Model simulations of attention to either eye during the remarkable saccade. Shown are remarkable saccade magnitudes when attention was focused on the aligned, unaligned eye or neither eye from two observers (PE & EV) (from Enright, 1992), and model performance.

Interestingly, Enright suggested that “attention given to the visual stimuli experienced by a given eye results in a greater “weighting” of those stimuli by the saccadic pulse generator”, which is consistent with changing the gain on the output of the conjugate component in our model.

To demonstrate the contributions of the conjugate and independent controllers to the remarkable saccade, we “lesioned” each class of controller by setting their inputs to zero (**Fig 9**). We then ran the model with parameters fixed to those used to fit to the raw traces of observer AB shown in Fig 5. When the conjugate controller is lesioned (left) the unaligned eye’s saccade reaches the target, and the aligned eye remains still. Therefore, removing the conjugate controller eliminates the remarkable saccade as expected. Lesioning the independent controllers (right) resulted in both eyes moving conjugately as might be expected. However, note that the saccades generated when the independent controllers are lesioned are of smaller amplitude than the unaligned eye saccade when these controllers are intact. This simulation result suggests that the independent controllers assist in guiding saccades to their final position.

**Fig 9.**
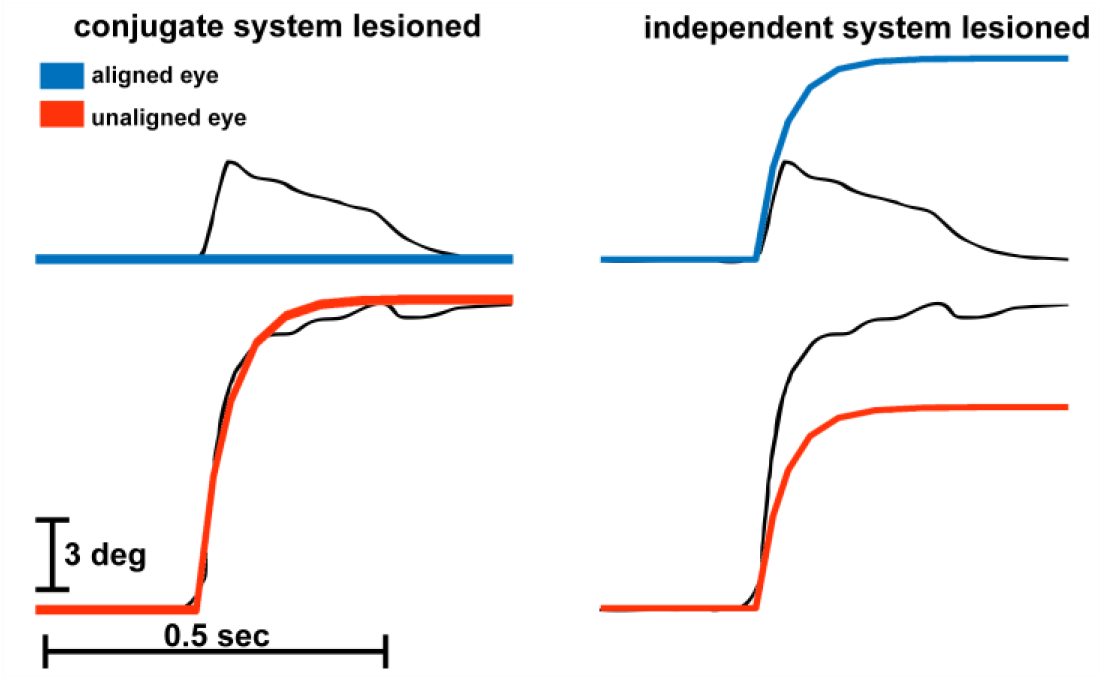
Model simulations when conjugate or independent systems are “lesioned.” Left, conjugate system lesion. Right, independent system lesion. Model traces (red/blue) are shown superimposed on raw traces from AB, Fig 5.

## Discussion

We demonstrate that a model proposed in Chandna et al. (2021) to explain our midline vergence pursuit results also generates inappropriate eye movements that occur during asymmetric vergence saccades known as “Remarkable Saccades” (Enright, 1992). Enright documented this curious phenomenon observed when using the Müller paradigm, where an unnecessary saccade from the aligned eye occurs that is in the same direction as that of the unaligned eye during gaze shifts between two targets at different depths. Our model replicates remarkable saccades for both convergent and divergent gaze shifts.

In Chandna et al. (2021), we occluded one eye and had observers pursue a target that moved smoothly back and forth on the midline. The covered eye showed little evidence of convergence or divergence. Instead, in most cases it rotated with a conjugate component.

Furthermore, the movement of the covered eye was desynchronized in time with that of the viewing eye, in violation of Hering’s Law. We proposed a conceptual model to explain these results (Chandna et al., 2021). Occluding one eye renders the retinal image motion created by the target’s midline motion ambiguous in terms of the target’s depth displacement. This is because the retinal image motion could correspond to an infinite family of trajectories either consistent with veridical motion of the target (purely moving in depth), moving on the frontoparallel axis, or moving on an axis between these extremes. How does the brain resolve this ambiguity? We proposed that it forms a “cognitive interpretation” of the target’s motion in depth using monocular depth cues and the memory of viewing the target’s motion binocularly. The cognitive interpretation of target motion in depth undergoes a neural conflict with well-established conjugate circuitry that resides in the brainstem. The signal resulting from the outcome of that conflict then drives the eyes.

Although not yet fully elaborated, we instantiated in our model a perceptual component on which the feedback loop is closed. We close the feedback loop on perception because previous work demonstrates that smooth pursuit follows perceived motion and not retinal motion. For example, when pursuing random dot stimuli (RDCs) the pursuit system minimizes the average dot velocity, and not that of an individual dot (Heinen & Watamaniuk, 1989). Interestingly, during pursuit of RDCs overall retinal slip can increase (**Fig 10**), and therefore some dots have *greater* retinal slip velocity than they would if an observer merely held their eyes stationary when the dot pattern moved.

**Fig 10.**
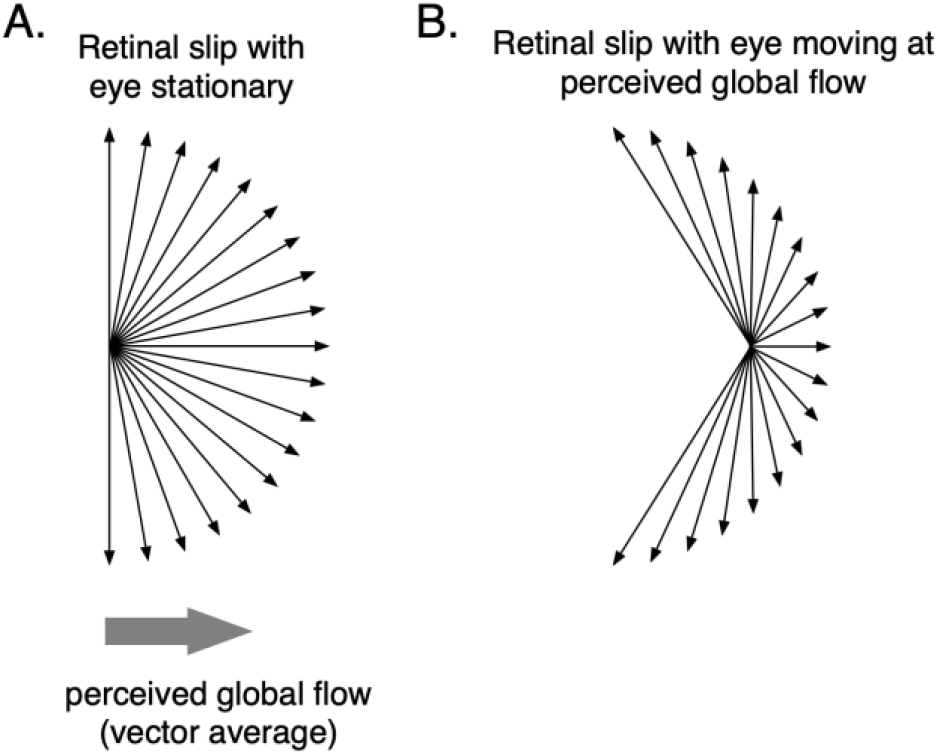
Retinal slip is not always minimized during pursuit. A) Retinal slip produced by dot movement of a random-dot cinematogram (RDC) (arrows) comprised of directions spanning 180° and the resulting perceived global motion (large shaded arrow). B) Remaining retinal slip produced by the RDC when the eye moves at the stimulus’ global velocity.

Other work also demonstrates that the pursuit system follows perception (Beutter & Stone, 1998; Madelain & Krauzlis, 2003), and not simply the motion on the retina as oculomotor models depict (e.g., Robinson et al., 1986; Krauzlis & Lisberger, 1989). The final component of the model is the virtual target, which incorporates the cognitive interpretation of the target, and it is used to rotate an eye without visual input, a component necessitated by Chanda et al. (2021) where an occluded eye continued to move despite lacking retinal input.

While based upon the Chandna et al. (2021) conceptual vergence model, this model also predicts several established oculomotor anomalies. One is the curiously slow vergence that occurs when saccades are executed to midline targets. Saccades to targets on a tangent screen have characteristic peak velocities that are proportional to their amplitudes, a relationship that is described by the “main sequence” (Bahill et al., 1975). However, when saccades are made to targets on the midline their peak velocities are much slower and no longer conform to the main sequence (Quinet et al., 2020). Greatly reduced eye velocity during vergence saccades is generally attributed to the vergence system being a sluggish system that rotates the eyes slower than does the conjugate system. Yet Chandna et al. (2021) casts doubt on the unitary vergence command upon which the classic vergence system relies. Our computational model generates sluggish midline vergence without the need for a separate vergence system. It suggests that sluggish vergence occurs because the eyes are oppositely directed, making the left and right conjugate controllers directly opposed, and mutual inhibition disables them. The slow controllers then operate in isolation to generate slow vergence rotations.

This manuscript was motivated by another even more surprising emergent property of the model, as it generates the remarkable saccades of asymmetric vergence (Enright, 1992). To the best of our knowledge, it is unknown how the brain generates remarkable saccades. Our model suggests they occur because the conjugate saccade generator triggers a saccade in both eyes, and the independent controller on the aligned eye, while damping its saccade, has insufficient potency to completely override it. This result highlights a key difference between our model and other existing models of oculomotor control; our model has separate inputs and outputs for each eye, while virtually all eye movement models are monocular at both the input and output (e.g., Robinson, 1973; Robinson et al., 1986; Krauzlis & Lisberger, 1989; Hung et al., 1986). It is the separate eye inputs and outputs that allow our model to generate remarkable saccades. This begs the question of why most previous eye movement models have not incorporated separate eye controls. We think this is likely because of the virtually universal acceptance of Hering’s Law. Acceptance of Hering’s law has led to most oculomotor scientists only recording one eye and assuming that the other eye behaves identically (or inversely), and hence behavior of both eyes are modeled as cyclopean. Notable exceptions are the independent control models (Helmholtz, 1867; King & Zhou, 2002). In addition, one prominent model although it has separate outputs does not use independent control (Zee et al., 1992) and its outputs incorporate Hering’s Law, which the Chanda et al. (2021) results challenge.

One aspect of the remarkable saccade emphasized by Enright (1992) is its within-subject variability. Our model does not capture remarkable-saccade variability because it, like virtually all oculomotor models which were designed to replicate the eyes’ averaged movement response, does not include internal noise. There are many places in our model where noise could and likely does arise including neural structures involved in detecting and encoding the visual stimulus (see Geisler, 1989) as well as structures that generate the visual percept upon which our feedback loop closes. It is generally accepted that our perceptions are not a perfect reflection of the physical stimulus. As such, some of the variability observed in Enright’s (1992) data is likely due to trial-to-trial variations in the percept as well as the ensuing motor commands based on those percepts. While a complete model of eye movements should incorporate sources of internal noise in order to capture eye movement variability, at this point ours does not.

Our model suggests a new conceptualization of the neural control of eye movements. We hypothesize that the basis of the conjugate component of our model resides in regions in the brainstem involved in generating saccades. The left and right components are theoretically subserved by burst neurons that reside in the superior colliculus (Sparks & Mays, 1980) and other regions of the brainstem such as the paramedian pontine reticular formation (PPRF) (Hepp & Henn, 1930). The architecture of the conjugate component is primitive and may have evolved from vestibular circuitry that drives the vestibuloocular reflex (VOR) which is found across numerous species, including fish. In support of this, while remarkable saccades have only been explicitly demonstrated in humans, we discovered evidence of it in a paper describing eye movements of zebrafish larvae (Bianco et al., 2011, Fig 5A). Furthermore, mutual inhibition between the left and right conjugate components in our model could be enacted in the brainstem through established mutual inhibition between inhibitory burst neurons (IBNs) (Shinoda et al., 2008).

We propose that the independent controllers of our model reside in cortex and may overlap circuitry that controls smooth pursuit, as smooth pursuit is more sluggish than saccades, and our model’s independent controllers are more sluggish than its conjugate controllers. Midline vergence is more sluggish than conjugate saccades, and it has been suggested that midline vergence and pursuit may be subserved by the same system (Erkelens et al., 1989). Furthermore, a smooth correction follows the rapid “saccade” during remarkable saccade execution (see Fig 3). If the regions involved in pursuit subserve the slow controllers, they might at least overlap areas in the middle temporal (MT) and the medial superior temporal cortex (MST) (Komatsu & Wurtz, 1988; Newsome et al., 1988). While MT and MST are in the motion system, and evidence is lacking that motion plays a role in asymmetric vergence or midline vergence saccades, to the best of our knowledge this has not been investigated. The interaction between signals from the independent controllers and the conjugate signal in our model could occur in the abducens nucleus, as neurons there carry conjugate signals arising from the PPRF, as well as activity corresponding to the rotation of an individual eye (Zhou & King, 1998).

Our model has implications for strabismus diagnosis and interventions. Most childhood strabismus is intermittent in nature (Birch et al., 1998; Nusz et al., 2006), and commonly manifests as Intermittent Exotropia (IXT) (Govindan et al., 2005). In cover tests of IXT patients, an outwardly deviated eye under cover returns to alignment with a saccade when the cover is removed. When that saccade is made, a simultaneous smaller saccade in the same direction often occurs in the fixating eye. This smaller saccade, like remarkable saccades, maladaptively displaces the fixating eye from the target. Consistent with the progression of remarkable saccades, the previously fixating eye subsequently recovers alignment with a slow, smooth movement (Economides, et al., 2017). Variations in this sequence including the frequency of occurrence and magnitude of the fixating eye’s saccade form the basis of clinical evaluation of IXT (Hatt et al., 2008). An explanation for the interplay between the two eyes has previously been a mystery as it is not explained by Hering’s Law. However, the HBC model offers a plausible explanation for the IXT cover-test result. In the framework of the model, the strength of the fixating eye’s independent control mechanism determines the frequency and magnitude of its displacement and return to the target in response to the deviated eye’s saccade - following removal of the cover. We hypothesize that the independent controller resides in cortex, possibly with input from motion processing regions in the brain. Our model suggests that the outcome of the cover test may reveal clues to the neural substrate underlying a patient’s IXT.

## Acknowledgements

Funded by NIH R01EY034626 and the Smith-Kettlewell Eye Research Institute.

